# Pharmacologic NLRP3 Inhibition Modulates Parkinson’s Disease-Associated Microglial Transcriptomic Signatures and Mitigates α-Synuclein–Triggered Neurodegeneration

**DOI:** 10.1101/2025.10.22.683837

**Authors:** Maria Luque, Magdalena Matic, Antonio Heras-Garvin, Jesus Amo-Aparicio, Kelvin C. Luk, Michaela Tanja Haindl, Michael Khalil, Damaris B. Skouras, Charles A. Dinarello, Nadia Stefanova

**Affiliations:** Laboratory for Translational Neurodegeneration Research, Division of Neurobiology, Department of Neurology, Medical University of Innsbruck, Austria; Department of Medicine, University of Colorado Denver, Aurora, CO, United States; Department of Pathology and Laboratory Medicine, University of Pennsylvania, Philadelphia, PA, United States; Department of Neurology, Medical University of Graz, Graz, Austria; Olatec Therapeutics Inc., New York, New York, United States

**Author notes:** the authors contributed equally (Maria Luque, Magdalena Matic, Antonio Heras-Garvin). corresponding author: Nadia Stefanova, Division of Neurobiology, Department of Neurology, Medical University of Innsbruck, Innrain 66/G3, 6020 Innsbruck, Austria.

**Keywords:** Parkinson’s disease, multiple system atrophy, NLRP3, neuroinflammation, α-synuclein, biomarker, disease-associated microglia, transcriptome, biomarker, therapy

## Abstract

**Background:** Parkinson’s disease (PD), the second most common neurodegenerative disorder after Alzheimer’s disease, and the rare disorder multiple system atrophy (MSA), are both characterized by intracellular accumulation of α-synuclein fibrils and early, sustained microglial reactivity in parallel to the neurodegeneration. Activation of the NLRP3 inflammasome in disease-associated reactive microglia is increasingly recognized as a key pathogenic driver and a promising therapeutic target in synucleinopathies. Dapansutrile (OLT1177®) is a selective, orally bioavailable NLRP3 inhibitor with a favorable safety profile in clinical trials for non-neurological indications. Here, we evaluated the therapeutic potential of dapansutrile in preclinical models of PD and MSA and explored the predictive and translational value of its effects.

**Methods:** Two established mouse models of synucleinopathy with nigral neurodegeneration were employed: the α-synuclein preformed fibril (PFF) propagation model and the transgenic PLP-α-syn model expressing human wild-type α-synuclein in oligodendrocytes. Pharmacokinetic analyses assessed plasma and brain exposure after oral administration. The efficacy of six-month dapansutrile treatment was examined in both preventive (post-PFF injection) and therapeutic (PLP-α-syn mice) paradigms, using behavioral, histopathological, and molecular readouts. Transcriptomic profiling of striatal and midbrain microglia identified differentially expressed genes (DEGs) associated with treatment and compared them with post-mortem transcriptomic signatures of disease-associated microglia in PD patients. Plasma IL-18 and neurofilament light chain (NfL) levels were evaluated as translational biomarkers.

**Results:** Chronic oral dapansutrile treatment at clinically relevant doses improved motor performance, reduced α-synuclein inclusions, attenuated gliosis, and mitigated nigral neurodegeneration in both models. Microglial transcriptomic analyses revealed that dapansutrile reversed key transcriptional signatures characteristic of PD-associated reactive microglia. Moreover, plasma IL-18 and NfL levels correlated with neuropathological and functional outcomes, supporting their potential as biomarkers of target engagement and treatment efficacy.

**Conclusions:** These data identify chronic NLRP3 activation as a shared and targetable mechanism in PD and MSA and highlight dapansutrile as a CNS-penetrant, clinically advanced candidate for disease modification in α-synucleinopathies. The observed transcriptomic reprogramming of microglia and the parallel changes in blood biomarkers provide a strong translational bridge to clinical development.

## BACKGROUND

Parkinson’s disease (PD) and multiple system atrophy (MSA) are progressive α-synucleinopathies that lack disease-modifying treatments. Although PD and MSA differ in histopathology and clinical course—Lewy body pathology in neurons in PD versus glial cytoplasmic inclusions in oligodendrocytes in MSA, differences in α-synuclein strains and in disease progression—both disorders share overlapping mechanisms and presentation that complicate early diagnosis and therapeutic intervention. [1–6] Chronic microglial activation and neuroinflammation emerge early in both conditions, paralleling α-synuclein aggregation and neurodegeneration, and highlight a potential common therapeutic target. [7–11].

The NLRP3 inflammasome, a multiprotein complex that drives caspase-1–dependent maturation of IL-1β and IL-18, is activated in disease-associated microglia in PD and MSA brains [12–14]. Experimental studies implicate NLRP3 activation in PD models [13, 15], but its role in α-synuclein–driven MSA pathology has not been defined. Addressing this gap is critical given the need for strategies that could target neuroinflammation across α-synucleinopathies.

Dapansutrile (OLT1177®) is an orally bioavailable, blood–brain barrier–penetrant, specific NLRP3 inhibitor currently in clinical development for non-neurological diseases. Its favorable safety, pharmacological profile and clinical advancement make it an attractive candidate for neurodegenerative disorders [16–18].

We show that dapansutrile improves motor function, reduces α-synuclein pathology, and limits neurodegeneration in both the PLP-α-syn transgenic model of MSA [19] and the α-synuclein preformed fibril (PFF) propagation model of PD [20]. Microglial transcriptomic profiling revealed that treatment reversed key disease-associated signatures characteristic of clinical PD described in single cell RNA sequencing studies [8, 9] underscoring the mechanistic basis of its efficacy. Together, these findings establish chronic NLRP3 activation as a common pathological driver of PD and MSA and highlight NLRP3 inhibition with dapansutrile—an oral, brain-penetrant drug already in clinical development—as a promising and immediately translatable therapeutic intervention for α-synucleinopathies.

## METHODS

### Study design

In our experiments, we applied two well-established synucleinopathy mouse models with nigral neurodegeneration: i) the PFF propagation mouse model [20] and ii) the transgenic PLP-a-syn mouse expressing human wild type a-synuclein in oligodendrocytes [21]. Sample sizes were determined by previous experience with these models. All in vivo experiments were performed according to the ARRIVE guidelines with the α of the Austrian Federal Ministry of Science and Research (BMFWF-2022-0.429.387).

In a first step, we aimed to provide a PK analysis of dapansutrile in the PLP-α-syn mouse model. Transgenic mice at 6 months of age received different doses of dapansutrile with drug-enriched food pellets for a period of 21 days. Plasma and brain levels of the drug were assessed at 2, 6, 8, 12, and 24h after ceasing the treatment.

Second, we aimed to assess the efficacy of six-month long oral treatment with dapansutrile (0g, 3.5g, and 7.5 g/kg feed pellets) initiated immediately after a stereotaxic α-synuclein PFF injection into the right striatum of C57BL/6 mice, applying behavioral and neuropathological outcomes of neuronal loss, synucleinopathy and neuroinflammation.

Finally, we aimed to assess the efficacy of six-month long oral treatment with dapansutrile (0g, 3.5g, and 7.5 g/kg feed pellets) initiated in six-month-old PLP-α-syn transgenic mice applying behavioral and neuropathological outcomes of neuronal loss, synucleinopathy and neuroinflammation. Transcriptomic analysis of striatal and midbrain microglia assessed the DEGs associated with the dapansutrile treatment and compared them to the clinical transcriptome signatures of disease-associated microglia in midbrain and profrontal cortex in PD patients [8, 9]. Plasma levels of IL-18 and neurofilament light chain (NfL) were measured as biomarkers of drug efficacy.

### Animals and treatment

We used two-month-old C57BL/6J and six-month-old PLP-α-syn mice [21] with equal sex representation and housed them under temperature-controlled pathogen-free conditions with a light/dark 12 hours cycle in the Animal Facility of the Medical University of Innsbruck.

Dapansutrile-enriched feed pellets (OLT1177®, Olatec Therapeutics LLC, New York, NY, USA) were provided at different concentrations (1.5g, 3.75g or 7.5g/kg pellets) selected based on previous studies in mice [22]. Mice had free access to either normal or drug-enriched food over specific periods as indicated in each experiment.

C57BL/6J PFF-injected mice were orally fed with normal feed, 3.75 g/kg and 7.5 g/kg dapansutrile-enriched feed pellets. Treatment was immediately started after PFF inoculation and lasted for a period of 6 months to assess treatment efficacy.

PLP-α-syn mice at 6 months of age received either normal diet or dapansutrile-enriched feed pellets (3.75g or 7.5g/kg) for the following 6 months to assess treatment efficacy.

### Pharmacokinetic studies of dapansutrile in PLP-α-syn mice

Six-month-old PLP-α-syn mice were randomized in four treatment groups and received either normal diet or dapansutrile-enriched feed pellets at 1.5g, 3.75g or 7.5g/kg for a period of 21 days. The feed delivery was ceased and plasma was collected after 2, 6, 8, 12, or 24 hours. We performed blood sampling from the facial vein, separated plasma by centrifugation at 5000 rpm for 10 min at 4°C and stored it at -80°C until further analysis. Mice were decapitated under deep isofluran anaesthesia, brains were immediately dissected, snap-frozen in liquid nitrogen and stored at -80°C until further analysis. Determination of dapansutrile in mouse plasma and brain homogenate samples was executed in accordance with established Standard Operating Procedures by Syneos Health (Princeton, NJ, USA) as previously [15]. Quantitation was based on the detection and integration of the product ion traces by mass spectrometry using MDS Sciex API6500 and Analyst® 1.7 (Applied Biosystems/MDS Sciex). Data were reported in ng/ml dapansutrile in plasma and ng/g dapansutrile in brain tissue.

### Stereotaxic surgery for the inoculation of PFFs in striatum

C57BL/6J mice at two months of age were randomized in four groups with equal representation of the sexes. Recombinant mouse monomeric α-synuclein and PFFs were generated, sonicated, characterized, and provided on dry ice by Dr. Kelvin Luk (for detailed protocols see [20]). The preparations were stored at -80°C and only thawed immediately before the stereotaxic injection. Stereotaxic injections were done under deep Isoflurane inhalation anesthesia on a stereotaxic frame (Kopf Instruments) using a 5µl Hamilton syringe. Monomeric α-synuclein or PFFs at concentration of 5mg/ml were injected in the right striatum with coordinates relative to Bregma (mm): anterior-posterior +0.2 mm, medial-lateral -2.0 mm, and dorsal-ventral -3.0 mm at a rate of 0.1 µl per minute (1µl total). The needle was left in place for at least 5 minutes after injection to minimize retrograde flow along the needle tract. Mice were placed on a heat-pad and monitored until complete recovery from anesthesia.

### Motor behavior analysis

#### Pole test

The test measures bradykinesia and coordination. Each mouse was placed on the top of a 50 cm high wooden pole. The time needed to turn downwards (T_turn_) and total time needed to climb down (T_total_) were recorded in five smooth runs. Mean values were used for the statistical analysis [23].

#### Challenging beam test

The test is applied to measure skillful movements when crossing a challenging beam detecting deficits in coordination, balance and motor performance. A horizontal 1-m-long beam elevated 20 cm above the surface, covered with a grid was employed. The beam’s width tapers from a wider starting point (3 cm) to a narrower endpoint (0.5 cm), increasing the challenge as the animal crosses the beam. Animals underwent habituation sessions for two consecutive days to familiarize with the beam and the goal box. This process reduces anxiety and ensures that performance during the test reflects motor abilities rather than novelty-induced stress. The animal was placed at the wider end of the beam facing the goal box and was allowed to traverse the beam towards the goal box containing the mouse home cage. If the animal did not initiate movement within a set time (e.g., 10 seconds), gentle encouragement was applied. Five performances were video-recorded per animal, and the number of slips per step of the hind limbs was measured. The best three performances per mouse were used for statistical analyses [23].

### Immunohistochemistry and immunofluorescence studies

PLP-α-syn mice were anesthetized with an overdose of thiopental and intracardially perfused with phosphate buffered saline (PBS pH 7.4, Sigma-Aldrich, Vienna, Austria). The right hemisphere was quickly dissected on ice and snap-frozen in liquid nitrogen for molecular analysis. The left hemisphere was fixed with 4% paraformaldehyde (PFA, pH 7.4, Sigma-Aldrich, Vienna, Austria) at 4°C overnight.

C57BL/6J mice receiving stereotaxic injections were sacrificed by intracardial perfusion with 4% PFA. Whole brains were post-fixed in 4% PFA at 4°C overnight. Fixed tissue was cryoprotected in 30% sucrose, slowly frozen in 2-methylbutane (Sigma-Aldrich, Vienna, Austria) and stored at -80°C until further use. 40-micron serial sections were cut using a cryostat (Leica Microsystems). Free-floating sections were stained using standard protocols. For immunohistochemistry, endogenous peroxidase was blocked by incubation in 0.3% H_2_O_2_ in PBS for 1 h. Sections were washed several times in PBS, and blocked with PBS containing 1% BSA, 5% normal serum, 0.3% Triton X-100 for 1 h at room temperature. Sections were then incubated with the respective primary antibodies (Table S12) overnight at 4°C and constant shaking. After 3 washes in PBS, sections were incubated with the respective biotynilated secondary antibodies (Table S12) for 1.5 h at room temperature. Following further 3 PBS washes, sections were incubated for 1 h at room temperature in ABC-complex (Vector Laboratories). Finally, the antibody binding was visualized by diaminobenzidine (Sigma). Sections were and mounted on slides and coverslipped with Entellan.

For immunofluorescence, the peroxidase block was skipped and the secondary antibodies used were Alexa-conjugated (Table S12). Slices were embedded in fluorescence embedding medium (Invitrogen) and kept in dark.

### Cytokine measurement in plasma and brain by ProcartaPlex® Multiplex Immunoassay

Tissue homogenization of striatum, cerebellum and midbrain was performed in RIPA-lysis buffer (Thermofisher, Waltham, MA, US) containing a mix of phosphatase and protease inhibitors (“phosSTOP” and “Complete, mini, EDTA-free,” Roche Applied Science). Samples were centrifuged at 16,000×g for 10 minutes at 4 °C and protein concentration was measured with BCA protein assay kit (Sigma-Aldrich, Vienna, Austria). Cytokines levels were simultaneously evaluated using a customized ProcartaPlex® Multiplex Immunoassay (Thermofisher, Waltham, MA USA) according to the manufacturer’s protocols. Each brain sample was loaded at a concentration of 2µg/µl. Plasma samples were loaded at 25 µl/well. Duplicates for each sample were measured and IL-1beta and IL-18 concentrations were defined as the mean value of the duplicates. Data are shown as pg/ml in plasma and pg/mg of total protein in brain homogenate.

### Neurofilament light chain (NfL) measurement in plasma by Single-Molecule Array (SIMOA)

Mouse plasma NfL concentration was quantified using commercially available NF-light™ V2 Advantage Kit (104037, Quanterix) with confirmed species cross-reactivity for mouse samples and analytical lower limit of quantification (LLOQ) of 0.345 pg/mL. Plasma samples were diluted 1:4 in the sample buffer as per manufacturer instructions. Single-molecule array (SIMOA) immunoassay was run on HD-X analyzer (Quanterix, Billerica, MA, USA) per manufacturer’s instructions for standards in technical triplicates and samples and controls in technical duplicates.

### Microglia isolation and characterization

Microglia were isolated from adult mouse striatum and midbrain using a protocol adapted from previously published methods with modifications [24, 25]. All steps were performed under consistently cold conditions to minimize ex vivo activation and transcriptional changes. Mice were deeply anesthetized with thiopental and perfused intracardially with 25 mL of ice-cold HBSS without Ca²⁺/Mg²⁺ (Thermo Fisher). Brains were quickly extracted, and the striatum and midbrain were dissected on ice in D-PBS (Thermo Fisher) supplemented with CaCl₂ (100 mg/L; Sigma-Aldrich), MgCl₂·6H₂O (100 mg/L; Sigma-Aldrich), D-glucose (1 g/L; Sigma-Aldrich), and sodium pyruvate (36 mg/L; Thermo Fisher). Tissue was dissociated using the Adult Brain Dissociation Kit (Miltenyi Biotec) following the manufacturer’s instructions, including processing in a gentleMACS Dissociator (Miltenyi Biotec), to generate single-cell suspensions. After dissociation, cells were spun down (500 × g, 4°C; Eppendorf), resuspended in D-PBS, and passed through pre-wetted 70 µm SmartStrainers (Miltenyi Biotec). After carefully removing the upper debris layers, cells were washed in D-PBS (1000 × g, 10 min, 4 °C), resuspended in PB buffer (D-PBS + 0.5% BSA), incubated with mouse CD11b MicroBeads UltraPure (Miltenyi Biotec) for 15 min at 4°C, and washed again. Prior to magnetic separation, an aliquot was taken for flow cytometry to determine the proportion of microglial cells in the sample. The labelled suspension was applied to MS columns (Miltenyi Biotec) in a magnetic separator, and magnetically bound cells were eluted with PB buffer. The eluate underwent a second round of magnetic separation to further increase purity. Isolated microglia were spun down (500 × g, 10 min, 4°C), snap-frozen in liquid nitrogen, and stored at −80°C until further processing. For flow cytometry, aliquots were stained with anti-CD11b-APC (clone M1/70, Thermo Fisher, 17-0112-83) and anti-CD45-FITC (clone 30-F11, Thermo Fisher, 11-0451-85) for 30 min at 4°C in the dark, washed, and resuspended in PB buffer prior to acquisition on a BD Accuri C6 (BD Biosciences) for characterization of the cell population by FACS using the markers CD11b and CD45 (Suppl Fig. 1).

### RNA processing and sequencing

Total RNA was extracted using the standard TRIzol method (Thermo Fisher), followed by purification with RNeasy Mini spin columns (Qiagen) and on-column treatment with RNase-free DNase (Qiagen), following the manufacturers’ instructions, and elution in DEPC-treated water. RNA quality and integrity were assessed using an Agilent TapeStation, with a mean RIN of 8.15 ± 0.73. Poly(A)-enriched libraries were prepared by Novogene UK and sequenced on an Illumina NovaSeq 6000 platform (150 bp paired-end), generating an average of 40 million reads per sample (6 GB raw data per sample).

RNA-seq raw reads were trimmed for residual adapter sequences and low quality sequences were removed using cutadapt [26]. RNA-seq reads were mapped to mouse and human reference genome (mm10 and hg38 respectively) with STAR aligner [27]. Read counts were obtained using featureCounts [28] and normalized using the normalization algorithms of DESeq2 [29]. Differential gene expression analysis of unterated versus dapansutrile-treated PLP-α-syn mice was performed in DESeq2 accounting for hidden batch effects by the removal of unwanted variation (RUVg, k4) method [30]. A threshold cut-off of P_adj_<0.1 (Benjamini-Hochberg procedure for false discovery rate (FDR) control) and │log_2_(FC)│>0.5 for higher biologically relevant sensitivity was applied.

### Pathway enrichment analysis

Applying the Gene Ontology (GO) terms for Biological Processes (BP) and Molecular Function (MF) as well as the Kyoto Encyclopedia of Genes and Genomes (KEGG), we performed enrichment analysis of differentially expressed genes with g:Profiler applying g:SCS multiple testing correction method with significance threshold of 0.05 to identify overrepresented terms in striatal and midbrain microglia of dapansutrile-treated versus untreated PLP-α-syn mice [31]. We performed separate enrichment analysis for the upregulated and downregulated genes either common for striatum and midbrain or specific for each structure. Selected enriched pathways were visualized by bar plots generated with SRplot [32].

### Cross-species comparison of microglial transcriptomic signatures of human Parkinson’s disease and effects of oral dapansutrile treatment in PLP-α-syn mice

We used single-cell RNA sequencing data on regulated genes in PD for disease-associated microglia located in the midbrain [9] and in the prefrontal cortex [8]. We mapped the human genes to orthologous mouse genes by applying g:Orth based on the information retrieved from the Ensembl database [31]. We excluded genes without determined orthologs or those with multiple orthologs from the analysis. To understand the relationships between the PD-regulated orthologs in human midbrain / human prefrontal cortex and the dapansutrile effects on the same microglial genes in the striatum and midbrain in the PLP-α-syn mouse, we applied descriptive hierarchical cluster analysis with complete linkage and Euclidean distance calculation using SRplot [32] on log2(FC) values, which were normalized for each gene and condition.

### Statistical analysis

Statistical analyses were performed using GraphPad Prism 10 (GraphPad). Depending on the type of data distribution, we applied the appropriate statistical test as indicated for each dataset. Differences with P < 0.05 were considered statistically significant. All data are presented as the mean ± standard deviation (SD) if not mentioned otherwise.

## RESULTS

### NLRP3 inflammasome activation in PD and MSA patients is replicated in α-synucleinopathy models

Extensive chronic microglial activation with upregulation of the NLRP3 inflammasome components occurs in the brains of PD and MSA patients [13, 14]. NLRP3 inflammasome activation has been reported across multiple PD models [13, 15, 33–36]. A previous study demonstrated that the α-synuclein PFF propagation model of PD induces inflammasome activation when compared with PBS-injected controls [13]. We extended this analysis by directly comparing PFF- and monomeric α-synuclein–injected mice. Immunohistochemistry unequivocally confirmed that microglial activation and the associated upregulation of ASC occurred in response to PFF α-synuclein (Fig. 1 A, B).

**Figure 1.**
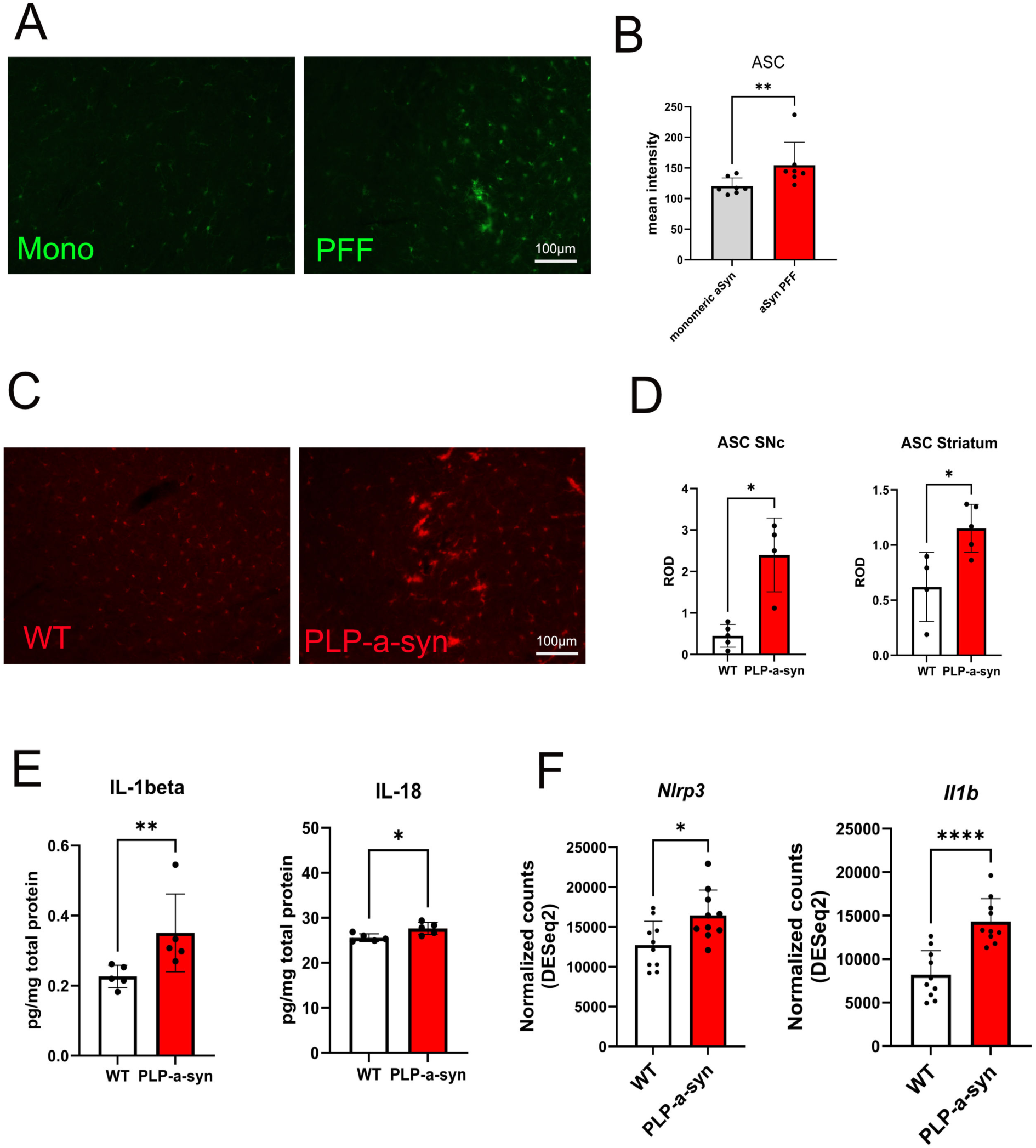
NLRP3 inflammasome activation in models of Parkinson’s disease (PD) and multiple system atrophy (MSA). (A) ASC immunofluorescence in the striatum of C57BL/6 mice 6 months after intrastriatal injection of monomeric α-synuclein (Mono) or α-synuclein preformed fibrils (PFFs). (B) Quantification of ASC fluorescence intensity (n = 7). (C) ASC immunofluorescence in brains of PLP–α-synuclein (PLP-α-syn) and wild-type (WT) mice. (D) Relative optical density of ASC immunofluorescence in the substantia nigra pars compacta (SNc) and striatum (n = 4–5). (E) Concentrations of interleukin-1β (IL-1β) and interleukin-18 (IL-18) (pg/mg total protein) in brain homogenates from WT and PLP-α-syn mice (n = 5). (F) Differential expression of *Nlrp3* and *Il1b* in microglia isolated from brains of WT and PLP-α-syn mice at 6 months of age (DESeq2 analysis; n = 10). Data are means ± SD. *P < 0.05, **P < 0.01, ****P < 0.0001 by unpaired two-tailed t test.

Microglial activation also characterizes MSA models with oligodendroglial α-synuclein inclusion pathology, including the PLP-α-syn transgenic mouse, in which human full-length wild-type human α-synuclein is expressed under the proteolipid protein (PLP) promoter in oligodendroglia [21]. In this MSA model, microglial activation progresses in a region-specific manner, paralleling striatonigral neurodegeneration and early cerebellar dysfunction [19], findings that are consistent with PET imaging results in MSA patients [37]. Immunohistochemistry analysis confirmed that at 6 months of age, at the time of initial microglial activation in the substantia nigra (SN) of PLP-α-syn transgenic mice, the adaptor protein ASC, a key component of the NLRP3 inflammasome, was significantly upregulated (Fig. 1C, D). In parallel, the downstream activation markers IL-1β and IL-18 were also significantly elevated in MSA mouse brains (Fig. 1E). Gene upregulation of *Nlrp3* and *Il1b* in the MSA mouse brain further supported these findings (Fig. 1F)

### Orally administered NLRP3 inhibitor dapansutrile is active in the CNS and blocks inflammasome activation in preclinical α-synucleinopathy

After confirming NLRP3 inflammasome activation in PD and MSA models with α-synuclein pathology, we tested whether dapansutrile could suppress this activation in the brain. In a primary experiment, we confirmed CNS penetration of dapansutrile following 21 days of drug-enriched feed in 6-month-old PLP-α-syn transgenic mice. The dose-dependent brain levels of dapansutrile two hours after food withdrawal correlated with plasma concentrations at the same time point (Fig. 2A-C).

**Figure 2.**
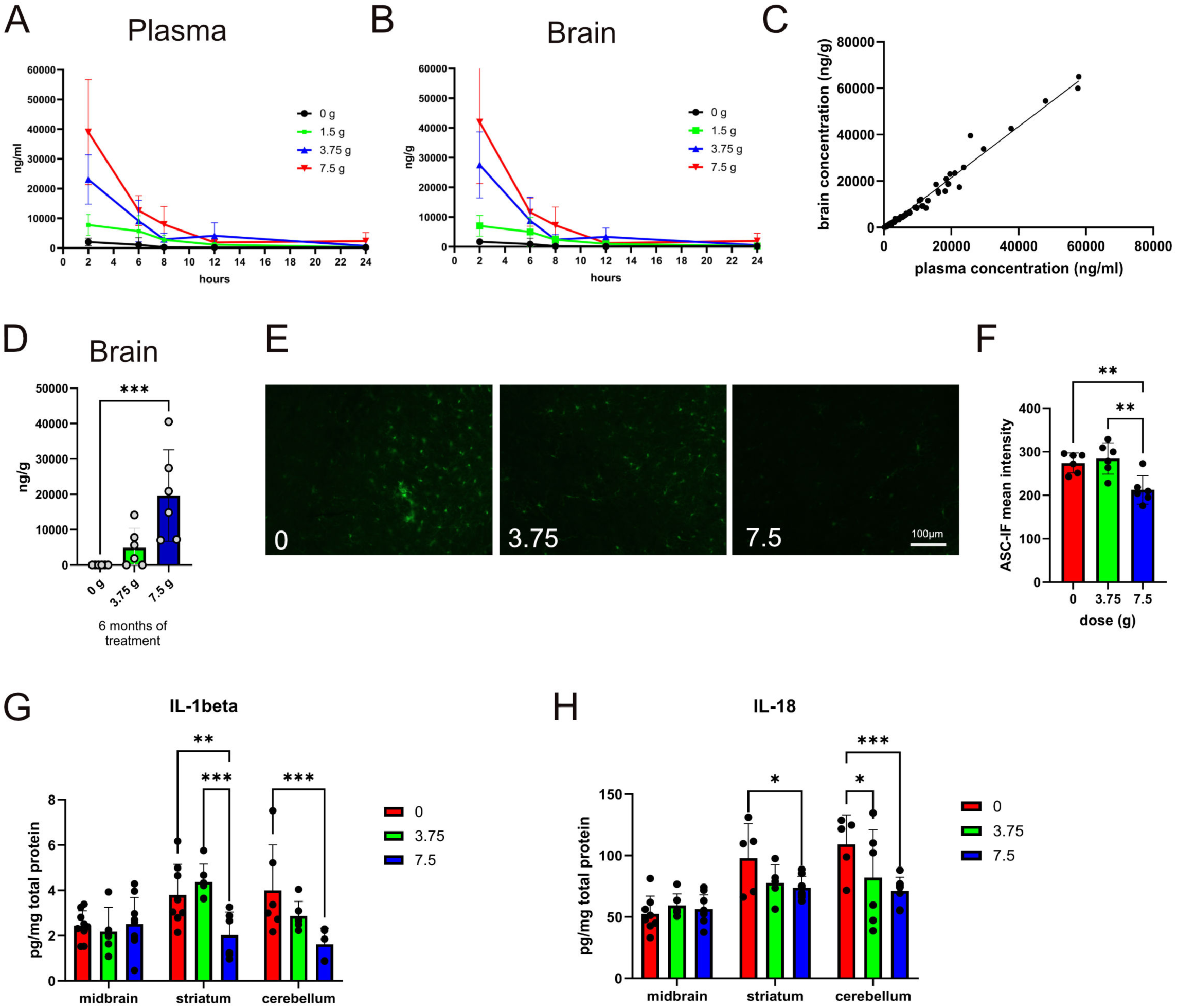
Dapansutrile crosses the blood–brain barrier (BBB) and suppresses NLRP3 activation in the brain. (A and B) Dose-dependent concentrations of dapansutrile in plasma (A) and brain tissue (B) of PLP–α-synuclein (PLP-α-syn) mice after 21 days of oral delivery in drug-enriched feed pellets (0, 1.5, 3.75, or 7.5 g/kg feed, with the 7.5 g/kg feed pellets equivalent to 1000 mg per day clinical dose), measured 2, 6, 8, 12, and 24 hours after treatment cessation (n = 6). (C) Correlation between plasma and brain concentrations (simple linear regression, R² = 0.9801, P < 0.0001). (D) Brain levels of dapansutrile in PLP-α-syn mice after 6 months of oral delivery (0, 3.75, or 7.5 g/kg feed; n = 6). (E) ASC immunofluorescence in brains of PLP-α-syn mice treated with different doses of dapansutrile. (F) Quantification of ASC fluorescence intensity (n = 6). (G and H) Duplex cytokine analysis of interleukin-1β (IL-1beta) and interleukin-18 (IL-18) levels in brain homogenates of PLP-α-syn mice treated with dapansutrile (0, 3.75, or 7.5 g/kg feed). Data are means ± SD. *P < 0.05, **P < 0.01, ***P < 0.001, by Kruskal–Wallis with Dunn’s post hoc test (D), one-way ANOVA with Tukey’s correction (F), two-way ANOVA with Fisher’s LSD test (G and H).

Next, we assessed brain concentrations of dapansutrile after 6 months of continuous drug-enriched feeding in PLP-α-syn transgenic mice, confirming sustained, dose-dependent CNS exposure (Fig. 2D). Dapansutrile treatment robustly suppressed NLRP3 inflammasome activation in the brain, as indicated by reduced ASC expression, along with decreased downstream IL-1β and IL-18 levels (Fig. 2E-H).

### Chronic oral dosing with dapansutrile protects against neurodegeneration induced by pathological α-synuclein aggregates in experimental PD and MSA mouse models

Dapansutrile bioavailability in the CNS after oral administration of 7.5 g/kg feed pellets achieved concentrations comparable to those shown to be clinically safe with 1000 mg dapansutrile per day in humans [16]. Therefore, we next studied whether oral dapansutrile confers neuroprotection and improves motor deficits in PD and MSA mouse models.

In the PD mouse model generated by intrastriatal injection of α-synuclein PFFs, we observed significant impairment in the challenging beam test and loss of dopaminergic neurons in the substantia nigra pars compacta (SNc) after 6 months, consistent with prior reports [20]. Oral dapansutrile treatment, initiated immediately after surgery, reduced slips per step on the beam and provided dose-dependent protection of nigral neurons (Fig. 3A–C).

**Figure 3.**
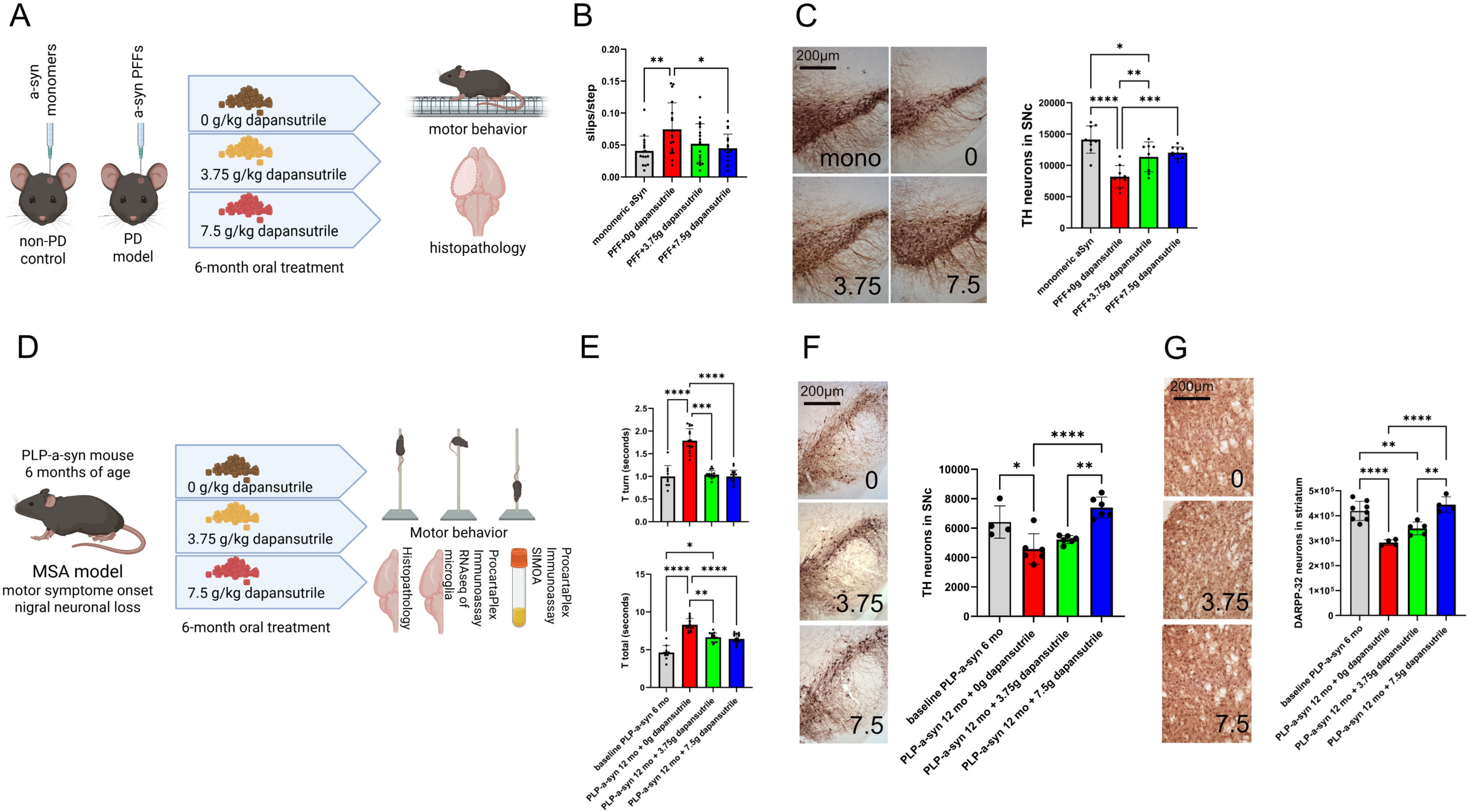
Dapansutrile improves motor performance and prevents neuronal loss in Parkinson’s disease (PD) and multiple system atrophy (MSA) mouse models. (A) Experimental design for PD modeling and preventive oral dapansutrile treatment with drug-enriched feed pellets for 6 months, followed by behavioral and histopathological analysis. (B) Slips per step in the challenging beam test (n = 16–19). (C) Tyrosine hydroxylase (TH) immunohistochemistry and stereological counts of dopaminergic neurons in the substantia nigra pars compacta (SNc) in PD and control mice (n = 8–10). (D) Experimental design for therapeutic 6-month oral dapansutrile treatment in PLP–α-synuclein (PLP-α-syn) mice starting at 6 months of age (MSA model with motor symptoms and early nigral degeneration), followed by behavioral, histopathological, biochemical, and microglial transcriptomic analyses, as well as plasma biomarker measurements [interleukin-18 (IL-18) for drug efficacy and neurofilament light chain (NfL) for disease progression]. (E) Pole test performance [time to turn (T_turn_) and total descent time (T_total_)] in PLP-α-syn mice at baseline (6 months) and after treatment (12 months) with dapansutrile (0, 3.75, or 7.5 g/kg feed; n = 10–17). (F) TH immunohistochemistry and stereological counts of dopaminergic neurons in the SNc of MSA mice (n = 4–6). (G) Dopamine- and cAMP-regulated neuronal phosphoprotein (DARPP-32) immunohistochemistry and stereological counts of medium spiny neurons in the striatum of MSA mice (n = 4–8). Data are means ± SD. *P < 0.05, **P < 0.01, ***P < 0.001, ****P < 0.0001 by one-way ANOVA with Dunnett’s multiple-comparison test (B), one-way ANOVA with Tukey’s post hoc test (C, F, G), or Kruskal–Wallis test with Dunn’s post hoc test (E).

To address whether dapansutrile may also modify symptom progression when initiated after disease onset, we treated 6-month-old PLP-α-syn mice, which already exhibited motor deficits, nigral neuronal loss, and showed progressive pathology in this α-synucleinopathy model [19]. Beam test detected no changes between 6 and 12 months of age and no changes linked to therapy (Fig. S1). Pole test analysis on untreated mice demonstrated significant worsening of T_turn_ and T_total_ in 12-month-old versus 6-month-old PLP-α-syn mice, accompanied by neuronal loss in both the SNc and striatum. Following oral dapansutrile treatment significantly improved motor performance and rescued dopaminergic nigral and medium spiny striatal neurons in a dose-dependent manner (Fig. 3 D–F).

### Pharmacological inhibition of NLRP3 inflammasome with dapansutrile suppresses neuroinflammation and lowers α-synuclein pathology in PD and MSA mice

Previous studies have shown that α-synuclein oligomers induce NLRP3 inflammasome activation, and that NLRP3 can affect the clearance of α-synuclein fibrils [38]. Conversely, pathological α-synuclein propagation is a central feature of α-synucleinopathies [39]. We examined α-synuclein propagation to the unilateral SNc 6 months after striatal PFF injection. Intracellular inclusions positive for hyperphosphorylated serine 129 α-synuclein (pS129) were detected in both neuronal somata and neurites, but not in mice injected with monomeric α-synuclein. Oral dapansutrile treatment significantly reduced nigral pS129 α-synuclein aggregates in the PFF PD model (Fig. 4 A–C).

**Figure 4.**
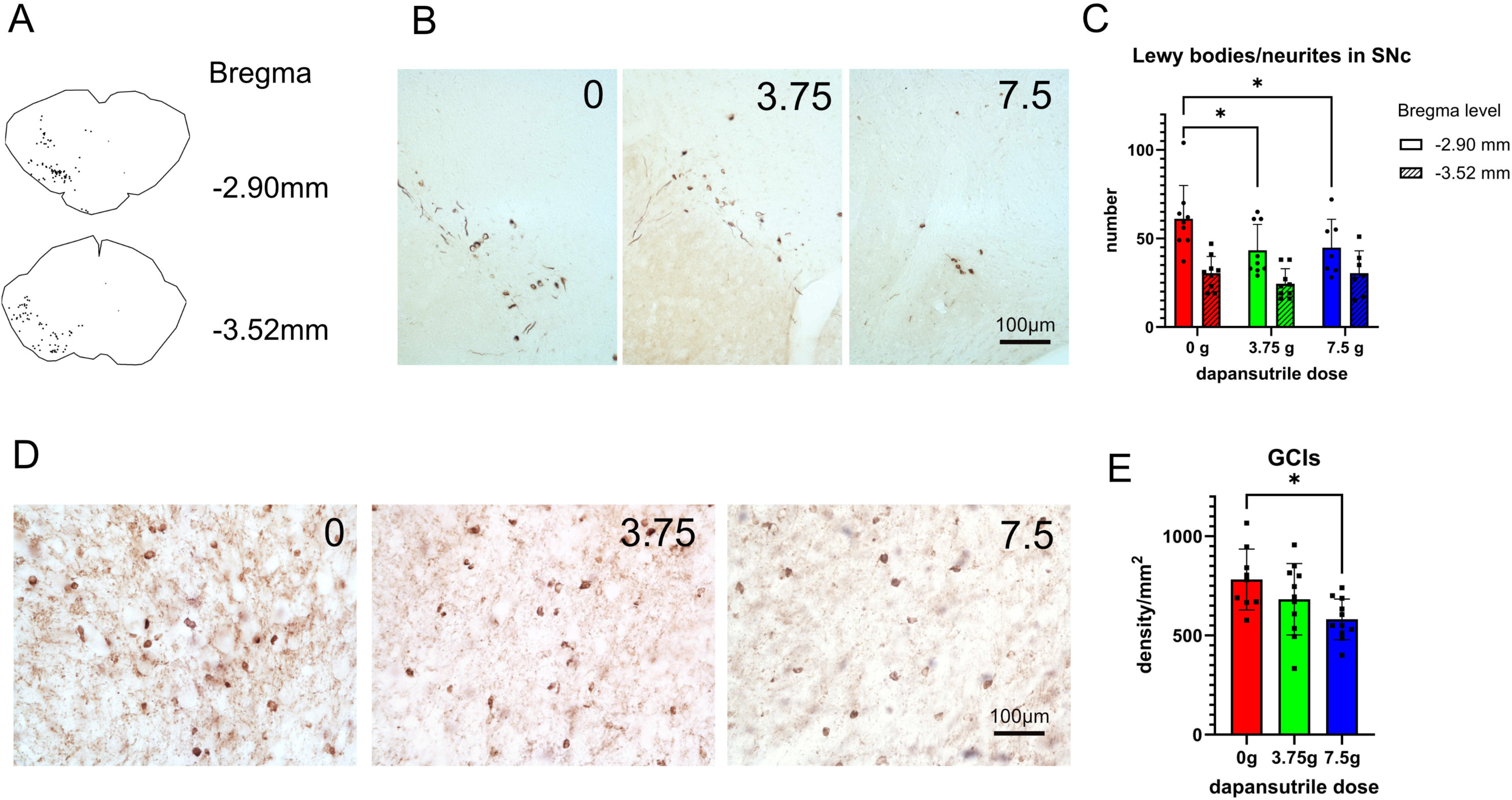
Dapansutrile reduces α-synuclein pathology in Parkinson’s disease (PD) and multiple system atrophy (MSA) mice. (A) Representative Lewy body distribution in the substantia nigra (SN) of wild-type (WT) mice 6 months after intrastriatal injection of α-synuclein preformed fibrils (PFFs) at two Bregma levels. (B) Immunohistochemistry for phosphorylated α-synuclein in PD mice. (C) Quantification of α-synuclein–positive inclusions in the substantia nigra pars compacta (SNc) of PD mice at two anatomical levels and the effect of oral dapansutrile treatment (n = 7–9). (D) Immunohistochemistry for phosphorylated α-synuclein in MSA mice. (E) Density of α-synuclein–positive glial cytoplasmic inclusions (GCIs) in MSA mice (n = 9–11). Data are means ± SD. *P < 0.05 by two-way repeated-measures ANOVA with Šídák’s multiple-comparison test (C) or Brown–Forsythe ANOVA with Dunnett’s T3 multiple-comparison test (E).

In the PLP-α-syn mouse model of MSA, α-synuclein inclusions accumulate in oligodendrocytes across multiple brain regions [19]. Dapansutrile treatment significantly reduced pS129 α-synuclein aggregates (Fig. 4D–F). Taken together, our findings implicate NLRP3 activation in the propagation and accumulation of pathological α-synuclein and demonstrate that oral dapansutrile mitigates this central disease process in α-synucleinopathies.

Astroglial and microglial activation have been documented in postmortem PD and MSA brains [40]. Pathological α-synuclein species elicit glial responses both in vitro and in vivo. In line with the presence of pS129 α-synuclein propagation and aggregation in PFF-injected and PLP-α-syn mice, we detected upregulation of key glial markers associated with neuroinflammation, which were significantly reduced after oral dapansutrile treatment (Fig. 5). This finding supports that dapansutrile, through NLRP3 inhibition, attenuates key downstream mechanisms of α-synucleinopathies.

**Figure 5.**
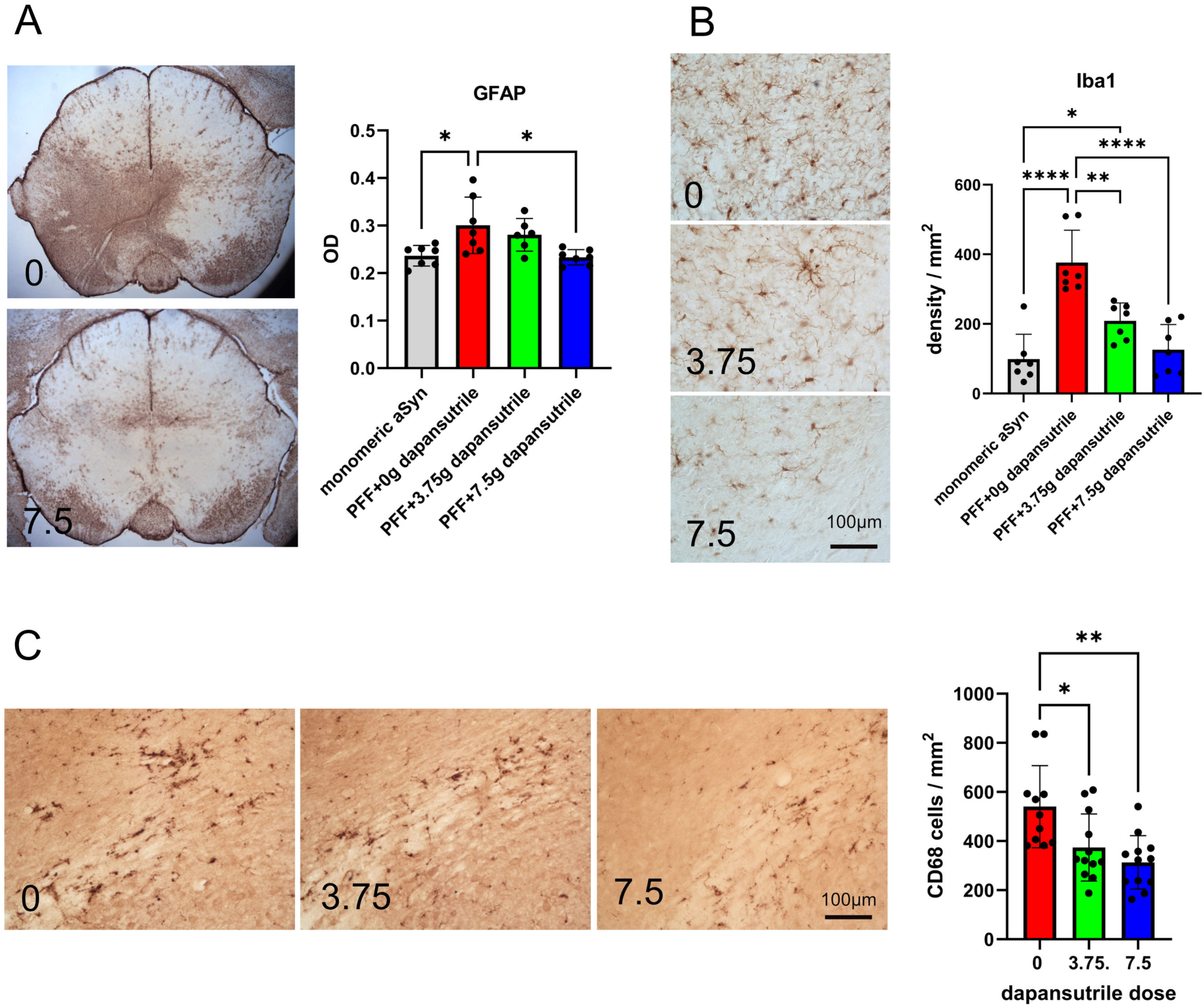
Dapansutrile reduces glial activation in Parkinson’s disease (PD) and multiple system atrophy (MSA) mice. (A) Glial fibrillary acidic protein (GFAP) immunohistochemistry and optical density of astroglia in PD mice (n = 6–7). (B) Ionized calcium-binding adapter molecule 1 (Iba1) immunohistochemistry and microglial density in PD mice (n = 7). (C) CD68 immunohistochemistry and density of phagocytic microglia in MSA mice (n = 11–12). Data are means ± SD. *P < 0.05, **P < 0.01, ***P < 0.0001 by Kruskal–Wallis test with Dunn’s multiple-comparison test (A) or one-way ANOVA with Tukey’s multiple-comparison test (B, C).

### Dapansutrile-induced transcriptional changes in microglia counteract disease-associated microglial profile in clinical PD

To characterize how dapansutrile modulates microglial activation in α-synucleinopathy, we performed transcriptome analysis of striatal and midbrain CD11b⁺ cells isolated from PLP-α-syn transgenic mice fed 7.5 g/kg dapansutrile-enriched or control pellets between 6 and 12 months of age. Flow cytometry of the isolated fraction confirmed a single CD11b⁺CD45⁺ population without a distinct CD45^high^ subset, indicating minimal peripheral macrophage contribution. Therefore, we refer to the isolated cells as microglia (Fig. 6A; Fig. S2). Next, we extracted RNA from the microglia, confirmed integrity (Table S1), and conducted RNA sequencing (RNA-seq) to identify the differentially expressed genes (DEGs) associated with treatment (Fig. 6B). A batch correction method was applied because samples were collected at different time points. Principal component analysis (Fig. S3) showed separation primarily by treatment group rather than batch.

**Figure 6.**
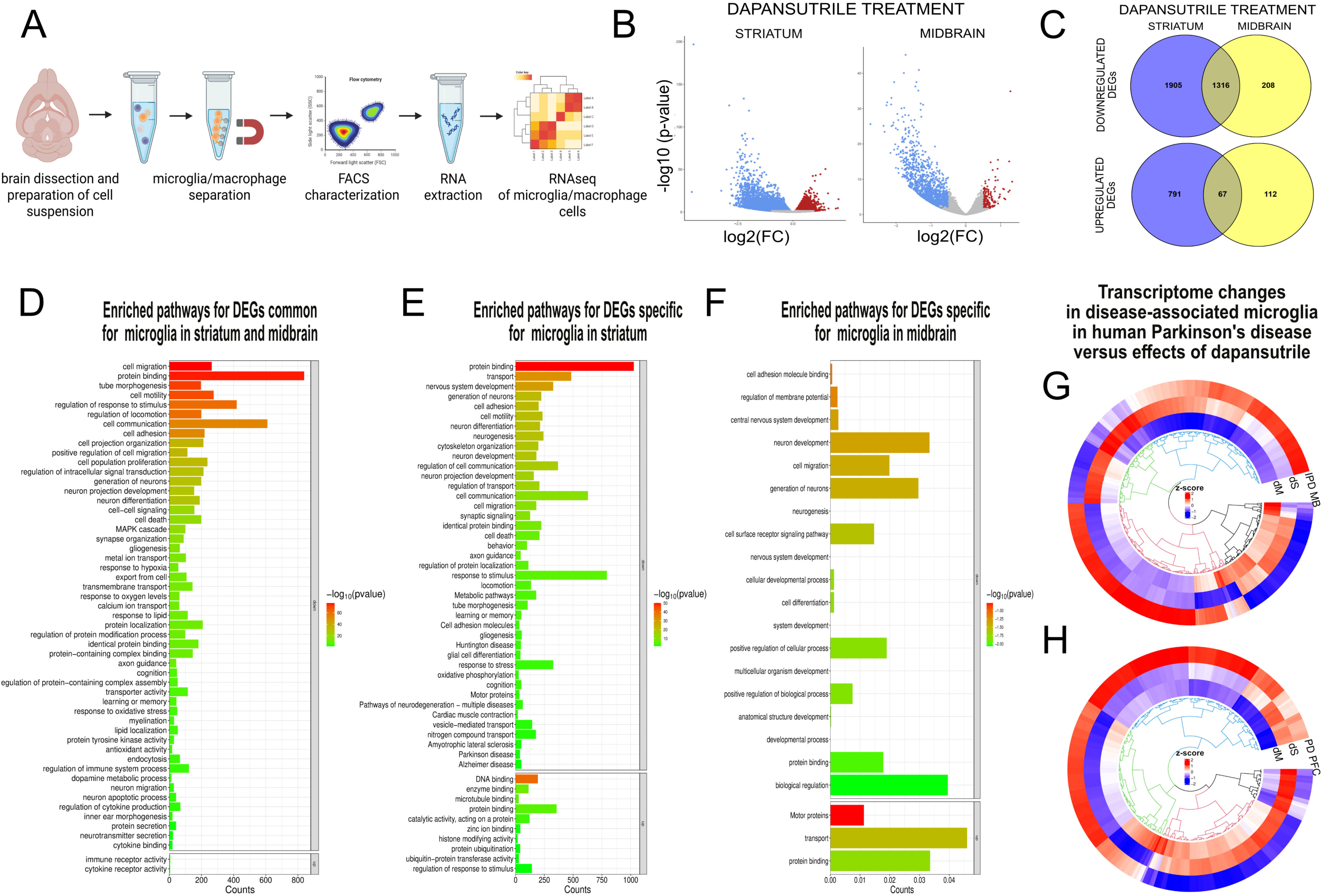
Transcriptomic effects of dapansutrile on microglia in PLP–α-synuclein (PLP-α-syn) mice and relevance to human Parkinson’s disease (PD). (A) Workflow of microglia isolation from adult PLP-α-syn mouse brains for RNA sequencing (RNA-seq). (B) Volcano plot of differentially expressed genes (DEGs) in the striatum and midbrain of 12-month-old PLP-α-syn mice after 6 months of oral dapansutrile treatment (7.5 g/kg feed; threshold P < 0.05, logFC > 0.5). (C) Venn diagrams of significantly up- and down-regulated microglial genes in the striatum and midbrain of PLP-α-syn mice after dapansutrile treatment. (D) Enriched pathways of overlapping downregulated and upregulated genes in striatal and midbrain microglia. (E) Enriched pathways of region-specific downregulated and upregulated genes in striatal microglia. (F) Enriched pathways of region-specific downregulated and upregulated genes in midbrain microglia. (G and H) Circular heatmaps with hierarchical clustering of orthologs dysregulated in post-mortem midbrain microglia from idiopathic Parkinson’s disease (IPD MB; in G) or prefrontal cortex microglia from PD (PD PFC; in H), compared with transcriptomic effects of dapansutrile in striatal (dS) and midbrain (dM) microglia from PLP-α-syn mice.

We identified 1,703 DEGs in midbrain microglia and 4,079 DEGs in striatal microglia after dapansutrile treatment (Table S2, S3). Most DEGs were downregulated in both regions, with a significant overlap of 1,316 downregulated genes (Fig. 6C). Pathway enrichment analysis revealed largely similar patterns across regions, despite differing numbers of DEGs. The significantly enriched pathways included regulation of microglial morphology and motility, immune responses, oxidative stress and hypoxia responses, neuronal processes (e.g., apoptosis, differentiation, projection and synapse development), and protein binding and modification (Fig. 6D; Tables S4, S5).

The striatum-specific DEGs were enriched for pathways linked to neuronal survival, neuron generation and differentiation, as well as neurodegeneration in multiple diseases including PD, Huntington’s disease, Alzheimer’s disease, locomotion, cognition and memory (Fig. 6E; Tables S6, S7), underscoring dapansutrile’s modulation of the ongoing neurodegenerative process in the striatum of PLP-α-syn mice, consistent with our histopathological findings. In the midbrain, pathway enrichment was less pronounced but in a similar direction, highlighting regulation of neuronal processes and protein metabolism (Fig. 6F; Tables S8, S9). Altogether, dapansutrile shifted the microglial profile toward reduced inflammatory morphology, migration, and neurotoxicity, while upregulating cascades involved in α-synuclein clearance and neuronal survival and connectivity.

Human and mouse microglial gene expression are broadly conserved, though species-specific differences exist [41]. To assess the clinical relevance of dapansutrile’s effects on microglial transcriptome, we applied two published single-cell RNA sequencing datasets from clinical PD. We mapped reported DEGs from PD midbrain [9] and PD prefrontal cortex [8] to mouse orthologs and compared their directionality with dapansutrile-induced changes (Fig. 6G; Tables S10, S11).

In both datasets, most PD-upregulated genes were downregulated by dapansutrile, while PD-downregulated genes were upregulated, supporting translational relevance. Interestingly, region-specific differences emerged, with clusters of PD-related genes showing differential dapansutrile responses in mouse midbrain versus striatum. This likely reflects regional specificity of microglial profiles, as dapansutrile’s effects in mouse midbrain aligned more closely with human PD midbrain (Fig. 6G) than PD prefrontal cortex (Fig. 6H). Conversely, dapansutrile’s effects in mouse striatum aligned better with PD prefrontal cortex, suggesting possible contributions of disease-stage activity in addition to regional specificity.

### Peripheral biomarkers of neuroinflammation and neurodegeneration track dapansutrile effects in experimental α-synucleinopathy

We assessed three plasma biomarkers – IL-1β, IL-18, and neurofilament light chain (NfL) – in untreated PLP-α-syn mice and mice receiving dapansutrile treatment. IL-1β and IL-18 were chosen as indicators of NLRP3 activity and target engagement, while NfL, a validated biomarker of neurodegeneration in PD and MSA [42–45], was used to test whether dapansutrile (7.5 g/kg feed) suppresses neurodegeneration in PLP-α-syn mice in a manner measurable by a peripheral biomarker.

IL-1β was undetectable in PLP-α-syn plasma using the ProcartaPlex assay and therefore unsuitable for analysis. IL-18 was reliably detected and showed a marked decrease with dapansutrile treatment (Fig. 7A). The absolute change in IL-18 plasma concentration during disease progression from 6 to 12 months of age in PLP-α-syn mice (Fig. 7B) correlated with the motor performance in the pole test (linear regression of IL-18 absolute change (pg/ml) to T_turn_: R^2^= 0.3149, P= 0.0019; and to T_total_: R^2^= 0.1826, P= 0.0233).

**Figure 7.**
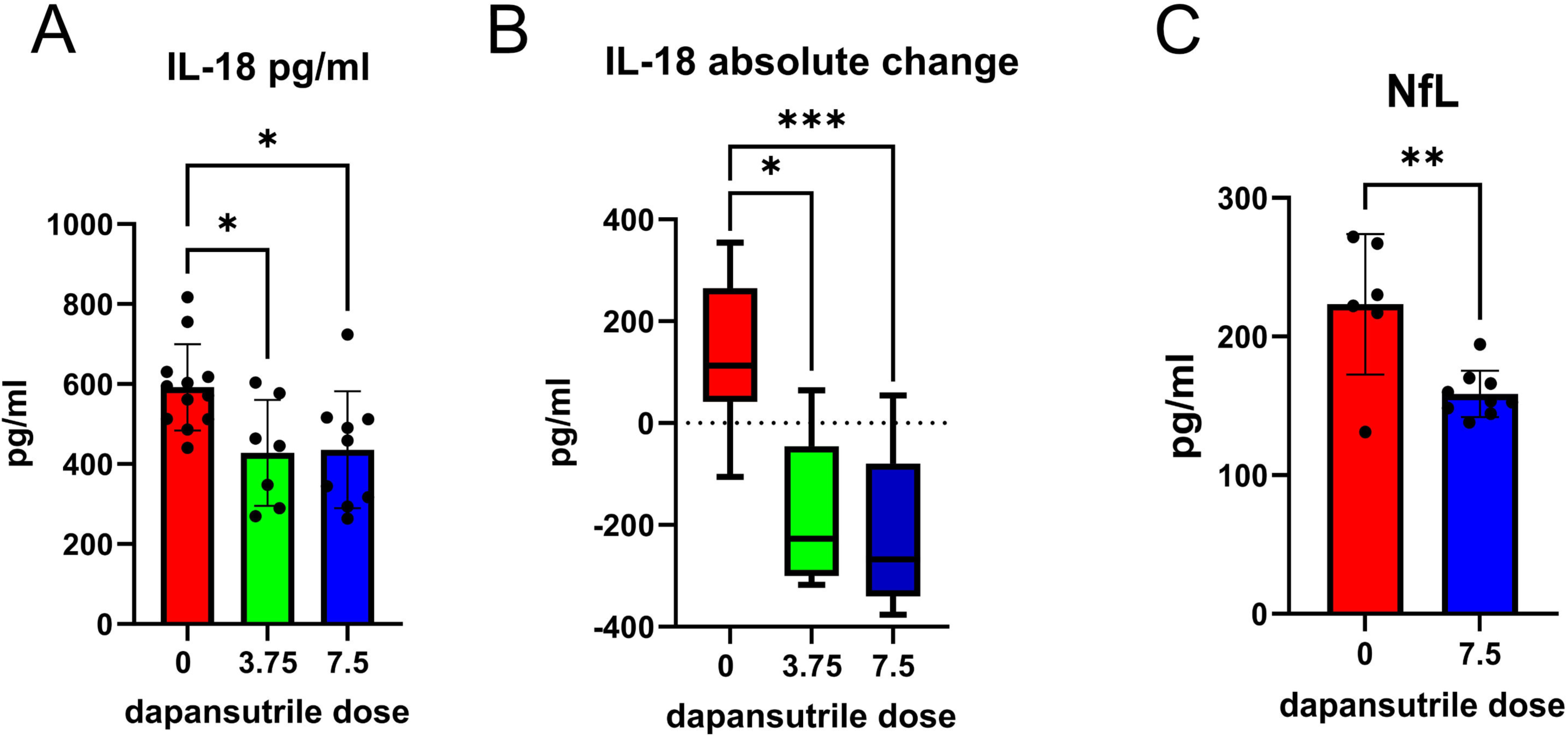
Plasma biomarkers of dapansutrile efficacy in Parkinson’s disease (PD) and multiple system atrophy (MSA) mice. (A) Plasma interleukin-18 (IL-18) concentrations measured by ProcartaPlex assay in PLP–α-synuclein (PLP-α-syn) mice treated with different doses of dapansutrile. (B) Absolute change in plasma IL-18 between 6 and 12 months of age in PLP-α-syn mice with or without dapansutrile treatment (n = 7–12). (C) Plasma neurofilament light chain (NfL) levels in PLP-α-syn mice with or without dapansutrile treatment (n = 6–9). Data are means ± SD. *P < 0.05, **P < 0.01, **P < 0.001 by Kruskal–Wallis test with Dunn’s multiple-comparison test (A, B) or Kolmogorov–Smirnov test (C).

Plasma NfL measured by SIMOA was significantly reduced with dapansutrile treatment of PLP-α-syn mice (Fig. 7C). Plasma NfL levels significantly correlated with pole test performance (linear regression of NfL (pg/ml) to T_turn_: R^2^= 0.3308, P= 0.0249; and to T_total_: R^2^= 0.3881, P= 0.0131). Taken together, these findings support IL-18 and NfL as peripheral biomarkers of dapansutrile efficacy, with direct relevance for monitoring treatment response in future clinical trials in PD and MSA.

## DISCUSSION

α-Synuclein aggregation is the hallmark pathology in α-synucleinopathies such as PD and MSA. The central role of misfolded α-synuclein in driving neurodegeneration and clinical deficits is well recognized, but multiple parallel mechanisms also contribute to disease progression. Designing therapeutic strategies that target several components of the pathogenic cascade remains a key goal for effective therapy in α-synucleinopathies.

Compelling clinical evidence for NLRP3 inflammasome activation in post-mortem PD and MSA brains and upregulation of key activation markers in microglia supports NLRP3 inflammasome as a promising disease-modifying target [13, 14]. NLRP3 inflammasome activation is linked not only to neurotoxic inflammatory responses but also to preventing the clearance of misfolded α-synuclein [38]. This double-edged action has made it an especially compelling target in α-synucleinopathies.

Initial studies in neurotoxin- and propagation-α-synuclein-induced models of PD suggested neuroprotection by NLRP3 inflammasome suppression [13, 15]. We provide strong evidence that dapansutrile, an oral selective NLRP3 inhibitor already in clinical development for metabolic, cardiovascular, and other inflammatory conditions [16, 17], is effective in α-synuclein–induced PD and MSA mouse models. Our findings demonstrate a dose-dependent effect of dapansutrile in two different synucleinopathy models, including modulation of glial responses, clearance of α-synuclein pathology, neuroprotection, and amelioration of motor deficits after symptom onset, thereby strongly supporting its immediate clinical translation in PD and MSA. Furthermore, our data provide microglial transcriptomic signatures of dapansutrile action that align with dysregulated microglial gene networks in human PD [8, 9], underscoring dapansutrile’s relevance to modulation of neuroinflammation, neurodegeneration, and proteostasis deficits linked to α-synucleinopathy.

The diagnosis and initiation of treatment in PD and MSA currently occur only after clinical symptom onset [3, 46]. This necessitates evaluating new interventions after symptom onset in preclinical models, in addition to preventive approaches used in prior NLRP3 studies [13, 15]. In our preclinical design, we applied side by side for the first time the α-synuclein PFF model and the PLP-α-syn mouse to assess preventive and therapeutic efficacy of dapansutrile. Our findings corroborated earlier observations in PD models (6-OHDA, MPTP, PFF) showing that NLRP3 targeting supports neuronal survival [13, 15]. The novelty and major impact of our study are based on the use of the PLP-α-syn MSA model, which enabled evaluation of dapansutrile after nigral neuron loss and early motor deficits had already emerged, providing a clinically relevant design. The PLP-α-syn mouse shows a well-established progression pattern: neuronal dysfunction and loss first occur in the lower brainstem and midbrain, later involving the striatum and cerebellum. Respectively, the motor phenotype begins with deficits on the beam test, correlating with nigral degeneration (similar to the PFF model), and progresses to pole test deficits associated with striatal degeneration [19, 47]. While beam test performance did not change within the test period (Fig.S1), dapansutrile treatment prevented further deterioration in the pole test, consistent with neuroprotection observed in the brain of PLP-α-syn mice (Fig. 3). This suggests that therapy may slow progression but does not reverse established disability. Our findings underscore the importance of early therapeutic initiation for successful outcomes in PD and MSA. This improved study design enhances the predictability and translational value of NLRP3 inhibitor efficacy for early-stage clinical trials.

We demonstrate that dapansutrile inhibits NLRP3, reducing IL-1β and IL-18 levels and ameliorating gliosis in both models (Figs. 2, 5), consistent with its mechanism of action. A major finding is the identification of dapansutrile’s transcriptomic signatures in the disease-associated microglia in synucleinopathy brains. By taking a conservative approach, we show that the transcriptional signatures of dapansutrile associate with modulation of multiple disease-associated DEGs linked to neuroinflammation, neurodegeneration, and proteostatic stress in both striatum and midbrain. Gene regulation was more extensive in striatum than in midbrain, consistent with advanced progression in the latter and more active degeneration in the striatum of PLP-α-syn mice between 6 and 12 months of age [19]. In both brain areas, we found disease-associated genes implicated by AD and PD GWAS, including *Apoe, App, Spp1, Lgals3*, and *Lrrk2* [48], significantly downregulated by dapansutrile.

Inflammatory mediators such as *Il1b* and *Tnf* were downregulated, while homeostatic genes such as *P2ry12* were upregulated [49]. The relevance of dapansutrile for clinical α-synucleinopathies is further underscored by dapansutrile-mediated downregulation of genes upregulated in human PD microglia as demonstrated by single-cell RNA-seq in postmortem midbrain [9] and prefrontal cortex [8]. Our cross-species analysis suggests that NLRP3 inhibition with dapansutrile modulates disease-associated microglial signatures relevant to clinical PD across two independent datasets. Our results propose that both regional specificity and disease stage may influence the effects of NLRP3 inhibition.

We reinforce the translational impact of our study by linking dapansutrile therapy to plasma biomarkers. We show that dapansutrile reduces plasma IL-18 (inflammation) and NfL (neurodegeneration), correlating with improved motor performance in PLP-α-syn mice. Together with the decreased brain levels of *Tspo* in striatum (Table S2), a widely used PET marker of microglial activation [50, 51], our findings suggest a feasible set of fluid and imaging biomarkers for future NLRP3 targeting trials in PD and MSA. Finally, we addressed the relationship between preclinical and clinical dosing. The more effective dose in mice that was concentrated into 7.5 g/kg feed pellets produced plasma concentrations comparable to those achieved with a safe 1000 mg/day dose in humans (Fig. 2A) [16, 17].

## Conclusions

Our study identifies chronic NLRP3 inflammasome activation as a shared and targetable pathogenic mechanism in PD and MSA. We provide compelling preclinical evidence that selective, oral NLRP3 inhibition with dapansutrile—a clinically advanced and CNS-penetrant small molecule—effectively attenuates α-synuclein pathology, neuroinflammation, and neurodegeneration across complementary models of synucleinopathy. By incorporating treatment initiation during established disease, transcriptomic profiling of microglial responses, and measurement of peripheral biomarkers, our study bridges critical gaps between preclinical and clinical investigation. The observed reversal of disease-associated microglial gene signatures toward homeostatic states, together with reductions in plasma IL-18 and NfL levels, supports the mechanistic and translational relevance of dapansutrile’s effects. Moreover, pharmacokinetic and dose–exposure analyses indicate favorable brain penetration and support safe clinical translation. We acknowledge that the experimental paradigms may not fully capture the presymptomatic course of human disease, which may influence treatment timing and long-term efficacy. Nonetheless, cross-species transcriptomic convergence and biomarker correlations provide the first predictive evidence for the biological efficacy of NLRP3 inhibition in α-synucleinopathies.

## LIST OF ABBREVIATIONS

6-OHDA: 6-hydroxydopamine
CNS: central nervous system
DEGs: differentially expressed genes
MPTP: 1-methyl-4-phenyl-1,2,3,6-tetrahydropyridine
MSA: multiple system atrophy
NfL: neurofilament light chain
NLRP3: NOD-, LRR-, and pyrin domain-containing protein 3
PD: Parkinson’s disease
PFF: pre-formed fibrils
PK: pharmacokinetic
PLP: proteolipid protein
pS129: hyperphosphorylated serine 129 α-synuclein
SNc: substantia nigra pars compacta
α-syn: α-synuclein

## Ethics approval and consent to participate

All in vivo experiments were performed according to the ARRIVE guidelines with the permission of the Austrian Federal Ministry of Science and Research (BMFWF-2022-0.429.387).

## Availability of data and materials

All data generated or analysed during this study are included in this published article and its supplementary information files.

## Competing interests

ML, declares no competing interests

MM, declares no competing interests

AHG, declares no competing interests

JAA, declares no competing interests

KCL, declares no competing interests

MTH, declares no competing interests

MK, has received travel funding and speaker honoraria from Bayer, Biogen, Novartis, Merck, Sanofi, Roche and Teva, serves on scientific advisory boards for Biogen, Bristol-Myers Squibb, Gilead, Merck, Neuraxpharm, Novartis, Alexion, Amgen and Roche and as a consultant for Roche. He received research grants from the Austrian MS Society, Biogen, Novartis and Roche.

DBS, Chief Executive Officer of Olatec Therapeutics

CAD, Chairman of Olatec’s Scientific Advisory Board, co-Chief Scientific Officer, receives compensation, and has equity in Olatec

NS, Director-at-large, Board of Directors and Chair of the Research Steering Council of Mission MSA, USA; Advisory board, Karl Golser Foundation, Italy; Advisory services for IONIS, Mitsubishi Pharma

## Funding

This study was supported by The Michael J. Fox Foundation for Parkinson’s Research (grant MJFF-022681). MM received a PhD fellowship by the intramural funding program of the Medical University Innsbruck PhD Research Training Groups, Project 2022-1-1.

## Authors’ contributions

ML has contributed to the stereotactic surgery, cytokine analysis and writing parts of the original draft.

MM has contributed to the behavioral analysis, tissue preparation and writing parts of the original draft.

AHG has contributed to the tissue preparation, microglia isolation with FACS analysis and writing parts of the original draft.

JAA has contributed to the conceptualization of the study.

KCL has contributed to the production and characterization of recombinant α-synuclein (monomeric and PFFs).

MTH has contributed to the SIMOA analysis of NfL.

MK has contributed to the SIMOA analysis of NfL.

DBS has contributed to the conceptualization of the study.

CAD has contributed to the conceptualization of the study.

NS has contributed to the conceptualization of the study, statistical and transcriptome analysis, supervision, funding acquisition, and writing parts of the original draft.

All authors read and approved the final manuscript.

## Acknowledgements

We thank Martina Wick for assistance in animal care and immunohistochemistry.

