## Supplementary material for "Pharmacologic NLRP3 Inhibition Modulates Parkinson’s Disease-Associated Microglial Transcriptomic Signatures and Mitigates α-Synuclein–Triggered Neurodegeneration": Fig. S1-S3

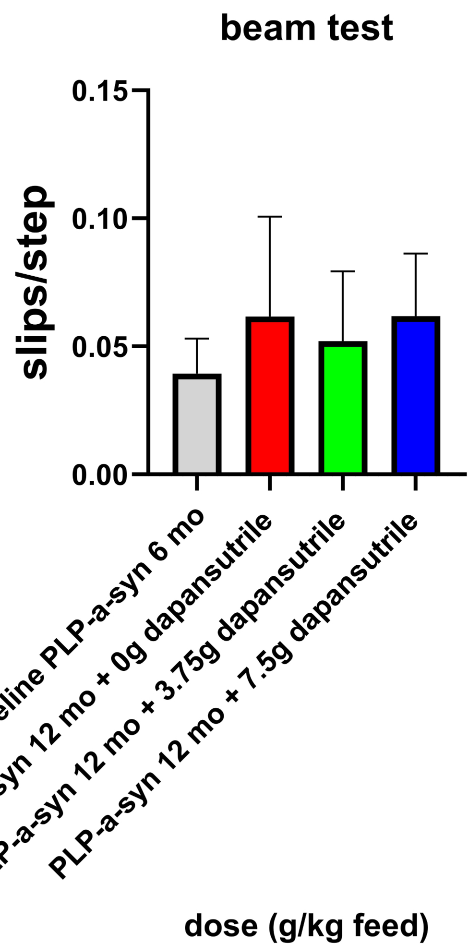


**Fig. S1.** Challenging beam test performance of PLP-a-syn mice at baseline (6 months of age) and at 12 months of age either untreated or treated with dapansutrile in two different doses.

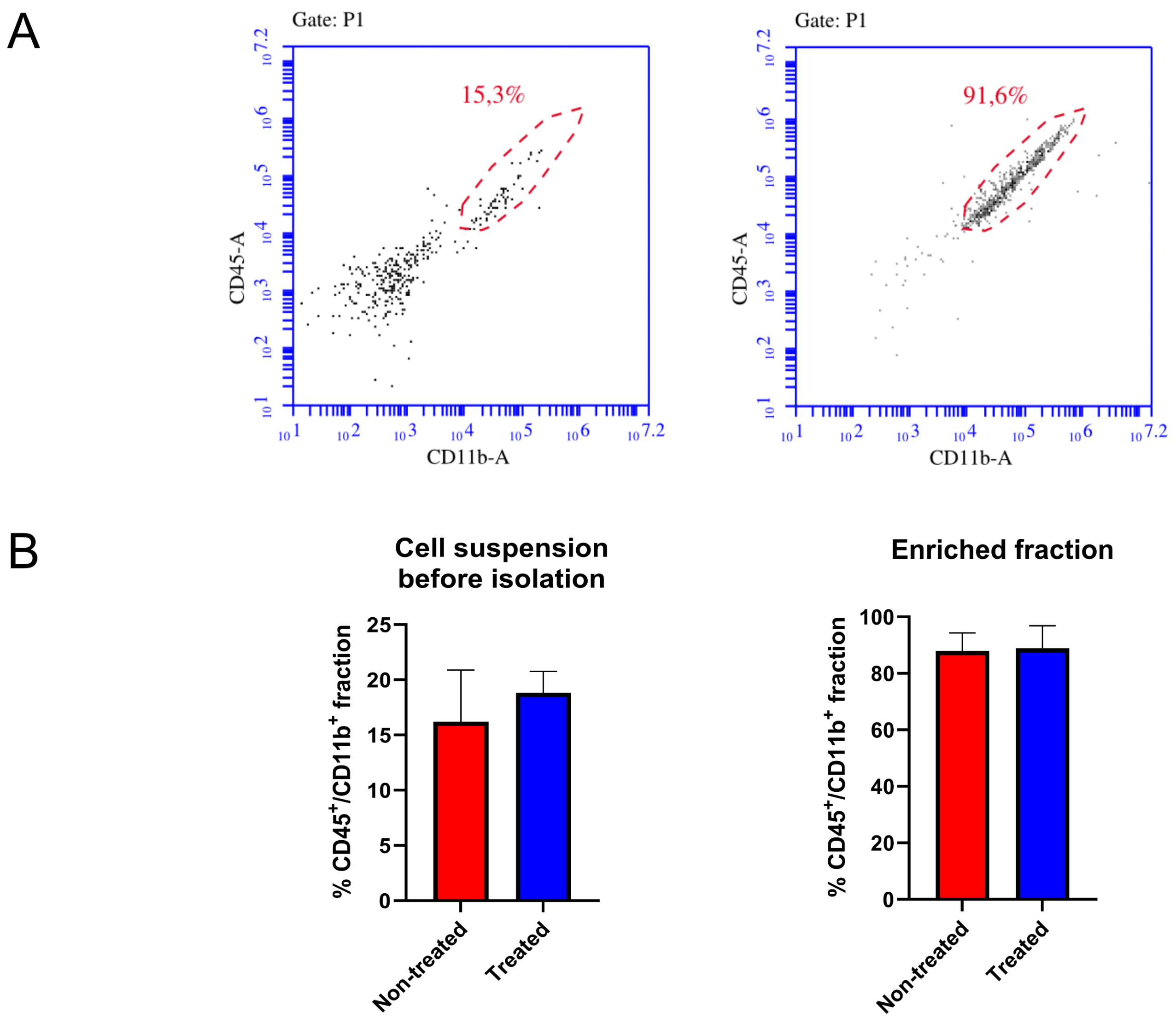


**Fig. S2**. Flow cytometry analysis. (A) Representative flow cytometry plots and (B) bar diagrams of percentage of CD45+/CD11b+ cells in total brain cell suspension and in the isolated enriched cell fraction used for RNAseq (n=5-6). Data shown as means ± SD.

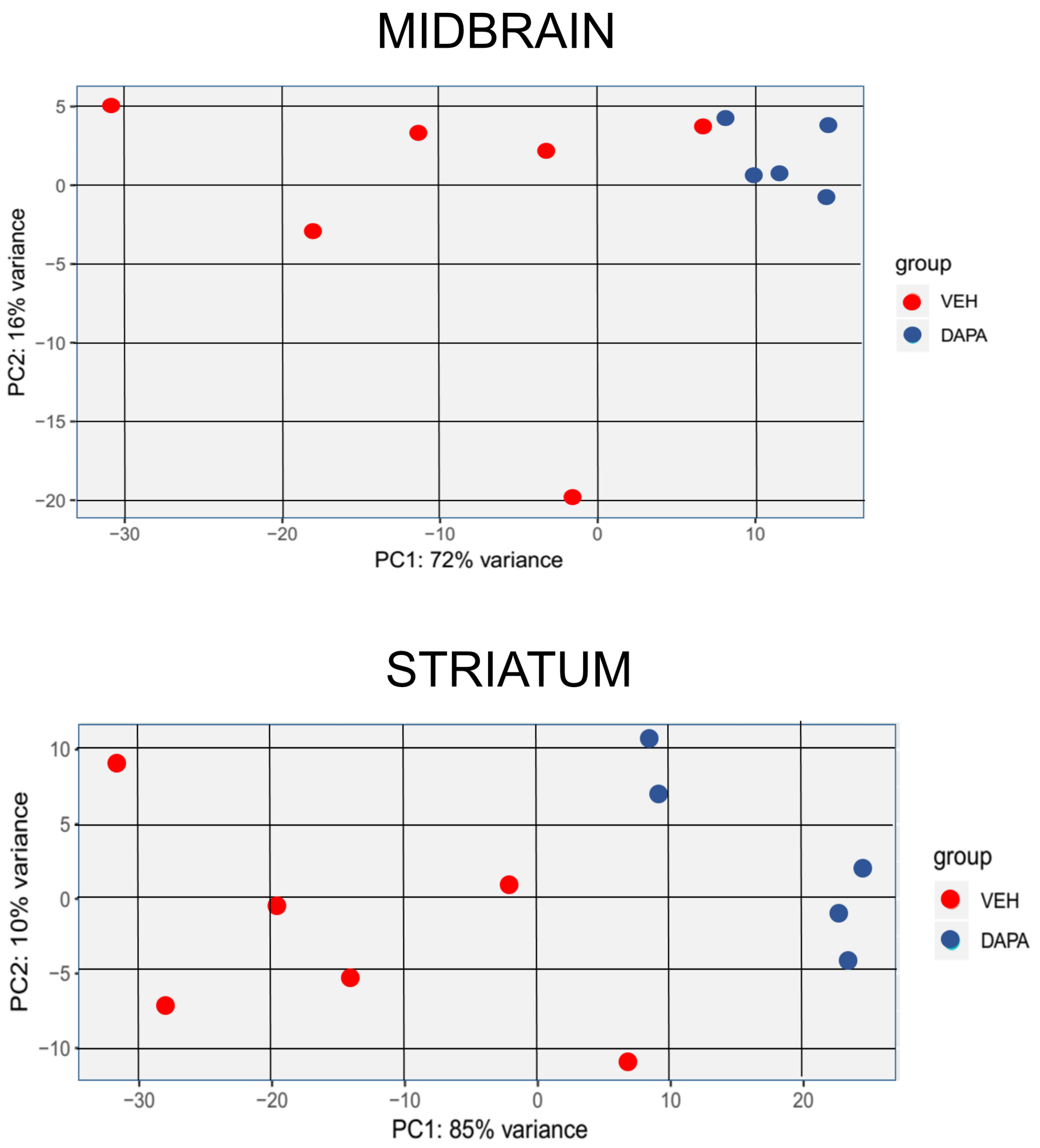


**Fig. S3**. Principle component analysis (PCA) plots of the raw data of RNAseq of microglia isolated from midbrain and striatum of PLP-a-syn mice, treated with vehicle (veh) or dapansutrile (dapa) as generated by DESeq2 analysis.
